# Investigating the Evolutionary Importance of Denisovan Introgressions in Papua New Guineans and Australians

**DOI:** 10.1101/022632

**Authors:** Ya Hu, Qiliang Ding, Yi Wang, Shuhua Xu, Yungang He, Minxian Wang, Jiucun Wang, Li Jin

**Affiliations:** State Key Laboratory of Genetic Engineering and Ministry of Education Key Laboratory of Contemporary Anthropology, School of Life Sciences, Fudan University, Shanghai 200438, China; CAS-MPG Partner Institute for Computational Biology, Shanghai Institutes for Biological Sciences, Chinese Academy of Sciences, Shanghai 200031, China; Department of Molecular Biology and Genetics, Cornell University, Ithaca, NY 14853, USA

**Keywords:** Denisovan, Archaic Introgression, Papua New Guinean, Australian, Local Adaptation

## Abstract

Previous research reported that Papua New Guineans (PNG) and Australians contain introgressions from Denisovans. Here we present a genome-wide analysis of Denisovan introgressions in PNG and Australians. We firstly developed a two-phase method to detect Denisovan introgressions from whole-genome sequencing data. This method has relatively high detection power (79.74%) and low false positive rate (2.44%) based on simulations. Using this method, we identified 1.34 Gb of Denisovan introgressions from sixteen PNG and four Australian genomes, in which we identified 38,877 Denisovan introgressive alleles (DIAs). We found that 78 Denisovan introgressions were under positive selection. Genes located in the 78 introgressions are related to evolutionarily important functions, such as spermatogenesis, fertilization, cold acclimation, circadian rhythm, development of brain, neural tube, face, and olfactory pit, immunity, etc. We also found that 121 DIAs are missense. Genes harboring the 121 missense DIAs are also related to evolutionarily important functions, such as female pregnancy, development of face, lung, heart, skin, nervous system, and male gonad, visual and smell perception, response to heat, pain, hypoxia, and UV, lipid transport, metabolism, blood coagulation, wound healing, aging, etc. Taken together, this study suggests that Denisovan introgressions in PNG and Australians are evolutionarily important, and may help PNG and Australians in local adaptation. In this study, we also proposed a method that could efficiently identify archaic hominin introgressions in modern non-African genomes.

## Introduction

Genomic introgressions from two extinct hominin species that used to reside on the Eurasian continent, namely Neanderthal and Denisovan, were detected in modern non-Africans (Green et al. 2010, Reich et al. 2010). Neanderthals were found to have admixed with the ancestors of all non-Africans. Denisovan introgressions, in contrast, were mostly found in PNG and Australians (Reich et al. 2011).

Despite the relatively low proportion, archaic hominin introgressions in non-Africans were found of great evolutionary importance. Neanderthal and Denisovan introgressions were found at immunity genes *STAT2*, *OAS* gene clusters, and *HLA* Class I genes (Abi-Rached et al. 2011, Mendez et al. 2012a, 2012b). Neanderthal introgression encompassing *HYAL2*, a gene related to the response to UV-B irradiation, was found at high frequency and under positive selection in East Asians (Ding et al. 2014a). Neanderthal introgression at the skin color gene *MC1R* was found to have introduced a functional missense allele V92M into the gene pool of anatomically modern human (AMH, Ding et al. 2014b). These studies suggest that archaic hominin introgressions might help modern non-Africans in local adaptation.

In addition, two previous studies have proposed methods in identifying Neanderthal introgressions in Eurasians at genome-wide scale (Sankararaman et al. 2014, Vernot and Akey 2014). These methods are based on the high divergence between Neanderthal introgressions and Africans, as well as the recent introgression time. The method proposed by Plagnol and Wall (2006), further extended by Vernot and Akey (2014), is based on high LD between variants absent from Africans. The method proposed by Sankararaman et al. (2014) is based on derived alleles that are consistent with Neanderthal but absent from Africans. They (Sankararaman et al. 2014, Vernot and Akey 2014) subsequently analyzed evolutionary importance of Neanderthal introgressions. To date, however, no study has focused on analyzing evolutionary importance of Denisovan introgressions in PNG and Australians.

One major issue in studying archaic hominin introgressions in AMH is to confidently demonstrate archaic origin of the identified introgressions. Improperly handling this issue could introduce false positives (Ding et al. 2014c). To address this issue, a previous study (Ding et al. 2014b) proposed a three-step approach. Putative introgressions should satisfy the following three criteria to be considered as from archaic introgression: (1) close phylogenetic relationship with Neanderthal or Denisovan; (2) the divergence time with Neanderthal or Denisovan postdates the African – archaic hominin population split time (i.e., 550 thousand years ago [KYA], Prüfer et al. 2014); and (3) reject the alternative model (i.e., incomplete lineage sorting model).

Here we present the analysis of Denisovan introgressions in PNG and Australians. We proposed a two-phase method to confidently identify Denisovan introgression. This method has relatively high detection power and low false positive rate. We then applied this method to twenty PNG and Australians genomes, and identified 1.34 Gb of Denisovan introgressive haplotypes and 38,877 DIAs. We found that 78 Denisovan introgressions were under positive selection, and 121 DIAs are missense. Genes in the 78 positively selected introgressions and harboring the 121 missense DIAs are related to a variety of evolutionarily important functions. In conclusion, Denisovan introgressions may play an important role in the evolution and local adaptation of PNG and Australians.

## Results

### A Two-phase Method in Identifying Denisovan Introgressions

Here we propose a two-phase method in identifying Denisovan introgressions in modern non-African genomes. The first phase is the *identification phase*. This method in identifying archaic hominin introgressions (denoted as *introgressive haplotypes* hereafter) is based on the long divergence time between archaic hominins and AMH. The second phase is the *filtering phase*, in which we employed an established three-step approach (Ding et al. 2014b) to reduce false positives.

Since the divergence time between African and archaic hominin (550 KYA, Prüfer et al. 2014) is much longer than that between African and non-African (<100 KYA, Jin and Su 2000), more mutations are expected to be accumulated on the archaic hominin lineage than on the non-African lineage after their divergence with Africans, and most of such mutations are absent from Africans. For two alleles on a SNP, if an allele is absent from all the African genomes, while could be observed on the non-African genomes, this allele is defined as an *E-allele* (Fig. 1). Such alleles have also been studied previously (Plagnol and Wall 2006, Wall et al. 2009, Sankararaman et al. 2014, Vernot and Akey 2014). There are three possible sources for E-alleles. The first source is mutations in archaic hominins that occurred after their divergence with Africans. The second source is mutations in non-Africans that occurred after their divergence with Africans. The third source is alleles that existed in the ancestors of Africans and non-Africans, but were not observed in African samples, due to either genetic drift or a limited African sample size. Based on simulation (Supplemental Text, Supplemental Table 1), we showed that introgressive haplotypes have higher density of E-alleles (denoted as *E-allele rate* hereafter) than the rest of the non-African genomes. We then implemented a hidden Markov model (HMM)-based approach with two hidden states (i.e., *lower* and *higher* E-allele rate) to partition each non-African genome into segments with different E-allele rates (see Material and Methods). The *first* hidden state includes segments with *lower* E-allele rates, and the *second* hidden state includes segments with *higher* E-allele rates. To reduce noises, segments with length less than 10 Kb will be discarded. We consider segments with higher E-allele rates (i.e., labeled as the second state) are more likely of archaic origin, and these segments will be subjected to filtering by the second phase to minimize false positives.

**Figure 1.**
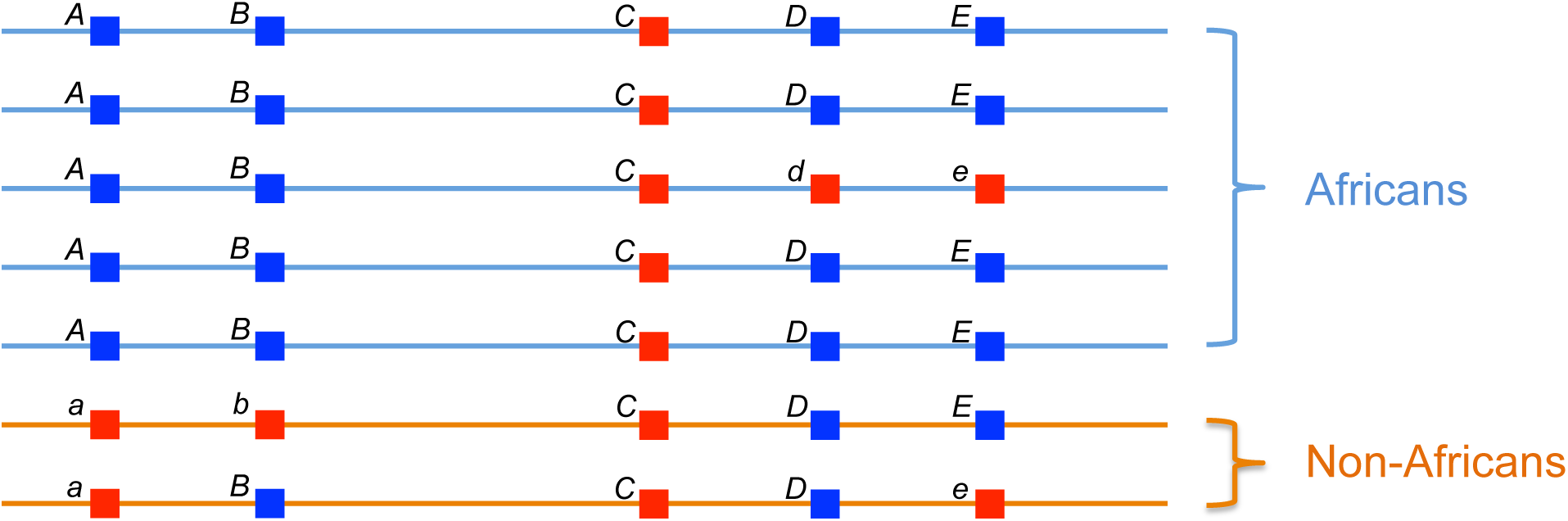
Definition of E-allele. For two alleles on a SNP, if an allele is absent from all the African genomes, while could be observed on the non-African genomes, this allele is defined as an E-allele. In the Fig. 1, alleles *a* and *b* are E-alleles, since both alleles are absent from all African genomes, while was observed in non-African genomes.

**Table 1.** Detection power and false positive rate of our two-phase method (λ from 6 to 15). See Supplemental Table 2 for full table (with λ from 1 to 25). PNG: Estimated using the demographic parameters of PNG. Eurasian: Estimated using the demographic parameters of Eurasian. Numbers in red are the maximum detection power and its corresponding false positive rate.

| $\lambda$ | PNG | | Eurasian | |
| --- | --- | --- | --- | --- |
|  | Detection Power | False Positive Rate | Detection Power | False Positive Rate |
| 6 | 0.7116 | 0.0201 | 0.8405 | 0.0604 |
| 7 | 0.7425 | 0.0210 | 0.8508 | 0.0606 |
| 8 | 0.7655 | 0.0207 | 0.8516 | 0.0624 |
| 9 | 0.7770 | 0.0213 | 0.8512 | 0.0631 |
| 10 | 0.7837 | 0.0219 | 0.8531 | 0.0644 |
| 11 | 0.7900 | 0.0224 | 0.8506 | 0.0643 |
| 12 | 0.7916 | 0.0233 | 0.8473 | 0.0651 |
| 13 | 0.7957 | 0.0241 | 0.8456 | 0.0672 |
| 14 | 0.7974 | 0.0244 | 0.8426 | 0.0675 |
| 15 | 0.7972 | 0.0250 | 0.8397 | 0.0677 |

The second phase is a filtering process designed to reduce false positive rate. We adopted a previously established three-step approach (Ding et al. 2014b) in the second phase. The segments identified by the first phase will need to satisfy the following three criteria to be considered as from Denisovan introgression: (1) the segment is phylogenetically closest to Denisovan, among Denisovan, Neanderthal, African, and chimpanzee sequences; (2) the divergence time between the segment and Denisovan is < 550 KYA (Prüfer et al. 2014), and the divergence time between the segment and Denisovan is less than that between the segment and Neanderthal; and (3) incomplete lineage sorting model is rejected for the segment by length of the segment (i.e., > 0.01089 cM, details see Material and Methods, Ding et al. 2014a). This filtering phase could reduce false positives introduced by ancient population structure, background selection, and GC-biased gene conversion (see Discussion).

In summary, we proposed a two-phase method in identifying Denisovan introgression in this section. We evaluated the detection power and false positive rate of this method in the following section.

### Detection Power and False Positive Rate of the Two-phase Method

We evaluated the performance (i.e., detection power and false positive rate) of the two-phase method by simulation. Demographic parameters were from Vernot and Akey (2014, for Eurasians, Africans and archaic hominins) and Prüfer et al. (2014, for PNG). The archaic hominin gene flow rate was from Vernot and Akey (2014), and the archaic hominin – African divergence time (550 KYA) was from Prüfer et al. (2014). Parameters for recombination hotspot and recombination rate were from Hellenthal et al. (2008).

Using msHOT (Hellenthal and Stephens 2007), we simulated non-African genomes with archaic introgressions. Sequence length for each simulation was set to 1 Mb. We define true archaic introgressions on the simulated non-African sequences as segments that (1) coalesced first with archaic hominin (based on the trees generated by msHOT); (2) diverged with archaic hominin < 550 KYA (based on the divergence times provided by msHOT); and (3) length of the segments are consistent with an introgression time after 550 KYA (i.e., genetic length > 0.01089 cM).

We then employed our method to the simulated non-African sequences, and evaluated the performance of our method. Detection power is defined as the proportion of the simulated archaic introgressions that were detected by the two-phase method, and false positive rate is defined as the proportion of the segments identified by the two-phase method that are not simulated archaic introgressions. We repeated the simulation for 1,000 times to evaluate detection power and false positive rate.

In the first phase of our method, there is an adjustable parameter named penalty value (λ). To obtain an optimized λ, we calculated detection power and false positive rate of our method using different λ values. Detection power reached maximum of 79.74% at λ=14, with false positive rate of 2.44% (Table 1, Supplemental Table 2). Therefore, in the following analyses on PNG and Australians, we set penalty value λ=14.

**Table 2.** A list of DIRs under positive selection and harbor evolutionarily important genes. For a full list of DIRs under positive selection, see Supplemental Table 4. Chr.: chromosome. Tag: position of the tag DIA of the DIR. Frequency: frequency of the Denisovan introgression. Top Position: position of the SNP with highest |iHS|. *r*^2^: LD *r*^2^ score between the tag DIA and the allele with highest |iHS|. All positions are in GRCh37. Genes in red are evolutionarily important.

| Chr. | Tag | Allele | Frequency | Highest \|iHS\| | Top Position | $r^2$ | Gene | Function |
| --- | --- | --- | --- | --- | --- | --- | --- | --- |
| 20 | 50305814 | T | 0.425 | 2.515 | 50316930 | 1.000 | *ATP9A SALL4* | Embryonic limb morphogenesis, neural tube closure |
| 5 | 35760424 | G | 0.400 | 2.015 | 35776369 | 1.000 | *SPEF2* | Spermatogenesis, fertilization, brain morphogenesis |
| 15 | 101432654 | G | 0.325 | 3.463 | 101449530 | 1.000 | *LOC100507452 ALDH1A3* | Face development, olfactory pit development |
| 2 | 168951543 | T | 0.275 | 2.552 | 168953002 | 1.000 | *STK39* | Response to stress, regulation of inflammation |
| 3 | 23612125 | C | 0.275 | 2.260 | 23630927 | 1.000 | *UBE2E2* | Cellular response to DNA damage stimulus |
| 15 | 66534679 | T | 0.225 | 3.648 | 66551937 | 1.000 | *MEGF11* | Retina layer formation |
| 3 | 64584252 | G | 0.225 | 2.294 | 64612943 | 0.861 | *ADAMTS9* | Positive regulation of melanocyte differentiation |
| 2 | 163726790 | A | 0.225 | 3.399 | 163665773 | 0.861 | *KCNH7* | Circadian rhythm |
| 3 | 179024584 | G | 0.200 | 2.758 | 179061537 | 1.000 | *ZNF639 MFN1 GNB4* | Negative regulation by host of viral transcription |
| 9 | 18942544 | T | 0.200 | 2.410 | 18969095 | 0.848 | *SAXO1* | Cold acclimation |

As aforementioned, we assume segments with *higher* E-allele rate identified by the first phase are more likely of archaic origin, and these segments will be subjected to filtering by the second phase. To provide support for this assumption, we hereby allow *all* segments identified by the first phase to enter the second phase, and re-calculated detection power and false positive rate (Supplemental Table 2). It was observed that the false positive rate rises, while maintaining similar detection power, which is in support of our assumption.

### Detecting Denisovan Introgressions in PNG and Australian Genomes

We applied the two-phase method to the sixteen PNG genomes and the four Australian genomes in the Simons Genome Diversity Project dataset with λ=14 where the detection power reached maximum in simulation. We identified 1.34 Gb of Denisovan introgressive haplotypes. We estimated that PNG and Australians carry 1.42±0.04% more Denisovan ancestry than Chinese Han in Beijing (CHB). This estimation is consistent with Meyer et al. (2012), which estimated that Papuan carry 2.0±0.9% more Denisovan ancestry than Han Chinese. We also replicated the findings in Mendez et al. (2012b), which reported Denisovan introgression encompassing the *OAS* gene cluster in Papuans. Furthermore, the identified Denisovan introgressive haplotypes diverged with Denisovan, Neanderthal, and Africans at 353.24 KYA, 641.97 KYA, and 957.77 KYA, respectively. This is consistent with reports in Mendez et al. (2012b) and Prüfer et al. (2014). The identified Denisovan introgressive haplotypes are included in Supplemental File 1, and is the data source for the subsequent analyses.

### Denisovan Introgressive Alleles

To investigate evolutionary importance of Denisovan introgressions, we identified alleles that were introduced by Denisovan introgression in PNG and Australians (denoted as *DIA* [Denisovan Introgressive Alleles] hereafter). Alleles that absent in Africans but carried by all Denisovan introgressive haplotypes were considered as *candidate* DIAs. To minimize effect of possible sequencing error, candidate DIAs that are singletons, doubletons, or tripletons were discarded.

We then implemented three stringent criteria identical to the three-step approach used in the second phase of the two-phase method to filter the candidate DIAs to minimize false positives. Specifically, a given candidate DIA could pass the stringent filter only if *all* of the haplotypes carrying the candidate DIA satisfy the three criteria: (1) phylogenetically closest to the Denisovan among PNG and Australian haplotypes, chimpanzee, African, Neanderthal, and Denisovan; (2) diverged with Denisovan < 550 KYA; and (3) genetic length is > 0.01089 cM to reject the incomplete lineage sorting model.

For example, for the candidate DIA rs117911431-T (chr2: 98206189, GRCh37), there are 80.00% (32/40) of the PNG and Australian haplotypes carrying this allele. To validate Denisovan origin for rs117911431-T, we first reconstructed a phylogenetic tree for PNG and Australian haplotypes, along with Denisovan, Neanderthal, African, and chimpanzee haplotypes (Fig. 2A). It was observed that all haplotypes carrying the rs117911431-T coalesced with Denisovan first, before joining the rest of haplotypes that do not carry this allele. Second, the haplotypes carrying the rs117911431-T diverged with Denisovan at 153.49 KYA, which postdates the divergence time between Denisovan and AMH (550 KYA). Third, the haplotypes carrying the rs117911431-T are 0.13 cM long, and therefore the incomplete lineage sorting model could be rejected (*p*<2.98×10^-16^). In summary, all haplotypes that carry the rs117911431-T satisfy the three stringent criteria, and thus the rs117911431-T is considered as a DIA.

**Figure 2.**
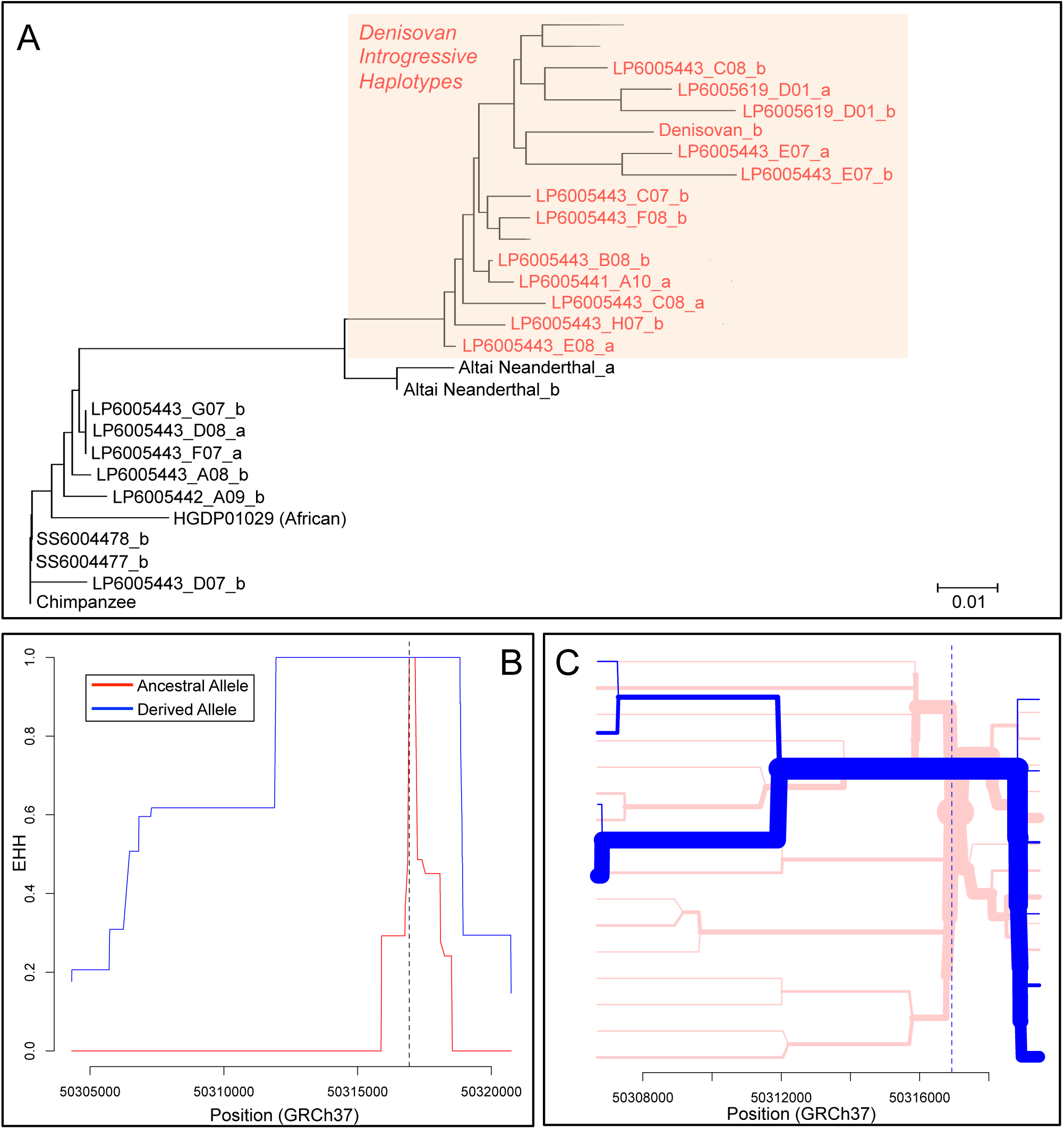
Phylogenetic tree, EHH analysis, and haplotype bifurcation graph. (A) Phylogenetic tree for PNG and Australian haplotypes, Denisovan, Neanderthal, and chimpanzee sequences, reconstructed using the neighbor-joining method in MEGA 6. Polymorphic sites within chr2: 98197888–98216073 (GRCh37, shared boundary of all rs117911431-T-carrying-haplotypes) were used to reconstruct the tree. Chimpanzee sequence was used as root. Trivial monophyletic clusters were collapsed to simplify tree presentation. Haplotypes carrying the rs117911431-T were colored in red. It was observed that these haplotypes coalesced with Denisovan haplotype first, before joining the rest of haplotypes that do not carry this allele, suggesting that the rs117911431-T is a DIA. (B) EHH graph of the SNP with highest |iHS| (rs372900695) in the DIR tagged by rs373805722-T. The derived allele of the SNP (rs372900695-A) is also a DIA, and is in complete LD with the tag DIA (rs373805722-T) of the DIR. The haplotype homozygosity for the derived allele (i.e., DIA) decays much slower than that of the ancestral allele, suggesting that the Denisovan introgression tagged by rs373805722-T is the target of positive selection. (C) Haplotype bifurcation graph of the SNP with highest |iHS| (rs372900695) in the DIR tagged by rs373805722-T. The haplotype homozygosity for the derived allele (i.e., DIA) decays much slower than that of the ancestral allele, which is consistent with Fig. 2B, and also suggests that the Denisovan introgression tagged by rs373805722-T is the target of positive selection.

In total, 38,877 alleles passed the aforementioned filtering process, and were thus considered as DIAs. To identify regions containing Denisovan introgression, we computed pairwise linkage disequilibrium (LD) *r*^*2*^ score for the identified DIAs. We define Denisovan introgressive regions (DIRs) as regions containing a set of DIAs in complete LD (*r*^*2*^=1.00), and a DIA randomly chosen from each DIR is used as a *tag DIA* to represent the DIR. In total, we identified 2,802 DIRs. The DIAs and DIRs are listed in Supplemental File 2 and Supplemental Table 3. It was observed that some tag DIAs are in LD (but not in complete LD), these tag DIAs could represent a same introgression event.

Among the 2,802 DIRs, frequency of Denisovan introgression in 2.64% (74/2,802) and 31.12% (872/2,802) of the DIRs reached 50% (20/40) and 25% (10/40), respectively.

**Table 3.**
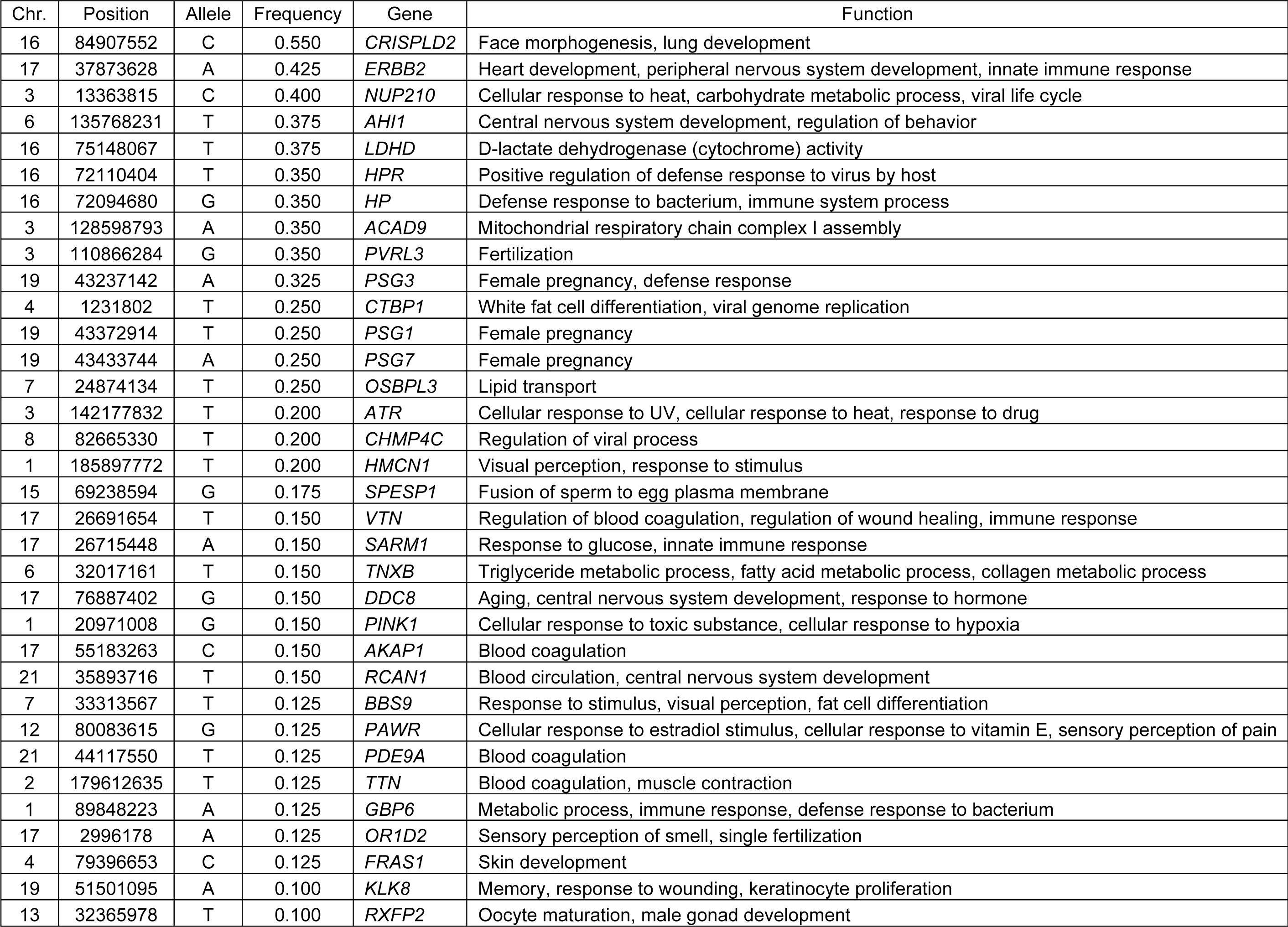
Missense DIAs harbored by evolutionarily important genes. Chr.: chromosome. See Supplemental Table 5 for a full list of missense DIAs.

| Chr. | Position | Allele | Frequency | Gene | Function |
| --- | --- | --- | --- | --- | --- |
| 16 | 84907552 | C | 0.550 | *CRISPLD2* | Face morphogenesis, lung development |
| 17 | 37873628 | A | 0.425 | *ERBB2* | Heart development, peripheral nervous system development, innate immune response |
| 3 | 13363815 | C | 0.400 | *NUP210* | Cellular response to heat, carbohydrate metabolic process, viral life cycle |
| 6 | 135768231 | T | 0.375 | *AHI1* | Central nervous system development, regulation of behavior |
| 16 | 75148067 | T | 0.375 | *LDHD* | D-lactate dehydrogenase (cytochrome) activity |
| 16 | 72110404 | T | 0.350 | *HPR* | Positive regulation of defense response to virus by host |
| 16 | 72094680 | G | 0.350 | *HP* | Defense response to bacterium, immune system process |
| 3 | 128598793 | A | 0.350 | *ACAD9* | Mitochondrial respiratory chain complex I assembly |
| 3 | 110866284 | G | 0.350 | *PVRL3* | Fertilization |
| 19 | 43237142 | A | 0.325 | *PSG3* | Female pregnancy, defense response |
| 4 | 1231802 | T | 0.250 | *CTBP1* | White fat cell differentiation, viral genome replication |
| 19 | 43372914 | T | 0.250 | *PSG1* | Female pregnancy |
| 19 | 43433744 | A | 0.250 | *PSG7* | Female pregnancy |
| 7 | 24874134 | T | 0.250 | *OSBPL3* | Lipid transport |
| 3 | 142177832 | T | 0.200 | *ATR* | Cellular response to UV, cellular response to heat, response to drug |
| 8 | 82665330 | T | 0.200 | *CHMP4C* | Regulation of viral process |
| 1 | 185897772 | T | 0.200 | *HMCN1* | Visual perception, response to stimulus |
| 15 | 69238594 | G | 0.175 | *SPESP1* | Fusion of sperm to egg plasma membrane |
| 17 | 26691654 | T | 0.150 | *VTN* | Regulation of blood coagulation, regulation of wound healing, immune response |
| 17 | 26715448 | A | 0.150 | *SARM1* | Response to glucose, innate immune response |
| 6 | 32017161 | T | 0.150 | *TNXB* | Triglyceride metabolic process, fatty acid metabolic process, collagen metabolic process |
| 17 | 76887402 | G | 0.150 | *DDC8* | Aging, central nervous system development, response to hormone |
| 1 | 20971008 | G | 0.150 | *PINK1* | Cellular response to toxic substance, cellular response to hypoxia |
| 17 | 55183263 | C | 0.150 | *AKAP1* | Blood coagulation |
| 21 | 35893716 | T | 0.150 | *RCAN1* | Blood circulation, central nervous system development |
| 7 | 33313567 | T | 0.125 | *BBS9* | Response to stimulus, visual perception, fat cell differentiation |
| 12 | 80083615 | G | 0.125 | *PAWR* | Cellular response to estradiol stimulus, cellular response to vitamin E, sensory perception of pain |
| 21 | 44117550 | T | 0.125 | *PDE9A* | Blood coagulation |
| 2 | 179612635 | T | 0.125 | *TTN* | Blood coagulation, muscle contraction |
| 1 | 89848223 | A | 0.125 | *GBP6* | Metabolic process, immune response, defense response to bacterium |
| 17 | 2996178 | A | 0.125 | *OR1D2* | Sensory perception of smell, single fertilization |
| 4 | 79396653 | C | 0.125 | *FRAS1* | Skin development |
| 19 | 51501095 | A | 0.100 | *KLK8* | Memory, response to wounding, keratinocyte proliferation |
| 13 | 32365978 | T | 0.100 | *RXFP2* | Oocyte maturation, male gonad development |

### Denisovan Introgressions in 78 DIRs were Under Positive Selection

We investigated whether the DIRs were under positive selection using the integrated haplotype score (iHS) test (Voight et al. 2006). Since the iHS is most effective in detecting positive selections with the frequency of selected sites ≥ 20%, we excluded DIRs with the frequency of the tag DIAs < 20% from the analysis for positive selection. Further, false positive rate of the iHS is not likely be elevated by archaic hominin introgression (unpublished results).

We first computed the standardized iHS for SNPs with minor allele frequency ≥ 5% in the twenty PNG and Australian genomes using the Whole-Genome Homozygosity Analysis and Mapping Machina (Voight et al. 2006). For each DIR, the Denisovan introgression in the DIR will be considered as under positive selection if more than one (not including one) of the SNPs in strong LD (*r*^*2*^≥0.80) with the tag DIA has standardized |iHS|≥2.00, since extreme iHS values (i.e., |iHS|≥2.00) caused by positive selection are usually in strong LD (Voight et al. 2006). We intend to reduce false positive by not including those regions with only one extreme iHS value. We further applied the Extended Haplotype Homozygosity (EHH) test (Sabeti et al. 2002) and drew haplotype bifurcation graph (Gautier and Vitalis, 2012) for the SNP with highest |iHS| to validate that the Denisovan introgression is the target of positive selection.

For example, for the DIR represented by rs373805722-T (chr20: 50305814, GRCh37, carried by 42.50% [17/40] of the PNG and Australian haplotypes), it was observed that three SNPs in complete LD with the rs373805722, namely rs372900695, rs373840702, and rs371966430, have standardized |iHS| of 2.515, 2.515, and 2.458, respectively. This observation suggests that the Denisovan introgression tagged by rs373805722-T could be under positive selection. This conclusion was further supported by EHH (Fig. 2B) and haplotype bifurcation graphs (Fig. 2C), since haplotypes carrying the DIAs have higher haplotype homozygosity. We thus concluded that the Denisovan introgression tagged by rs373805722-T was under positive selection in PNG and Australians.

Totally, we found that Denisovan introgressions in 78 DIRs were under positive selection (Table 2, Supplemental Table 4). Further, genes located in the 78 positively selected DIRs are related to a variety of evolutionarily important functions, such as reproduction (*SPEF2*), response to environmental conditions (*SAXO1* [temperature], *STK39* [stress], *UBE2E2* [DNA damage], and *KCNH7* [circadian rhythm]), development (*SALL4* [limb and neural tube], *SPEF2* [brain], *ALDH1A3* [face and olfactory pit], *MEGF11* [retina], and *ADAMTS9* [melanocyte]), and immunity (*STK39* and *ZNF639*). Positive selection on the DIRs harboring these genes may suggest that Denisovan introgressions at these genes are adaptive in PNG and Australians.

### One hundred and twenty-one DIAs are Missense Alleles

Besides positive selection, we are also interested in missense DIAs, since missense alleles are more likely to be functional. We obtained annotation for all 38,877 DIAs from the Variant Effect Predictor of the Ensembl Genome Browser (Flicek et al. 2014). Among all 38,877 DIAs, we found 121 missense DIAs (Table 3, Supplemental Table 5).

Similar to the above discovery that the genes located in the positively selected DIRs are evolutionarily important, genes harboring the 121 missense DIAs are also of evolutionary importance. Functions of the genes harboring the missense DIAs include reproduction (*PVRL3*, *PSG1*, *PSG3*, *PSG7, SPESP1, OR1D2*, and *RXFP2*), response to environmental conditions (*NUP210* [heat], *ATR* [heat, UV, and drug], *PAWR* [vitamin E and pain], *OR1D2* [smell], *HMCN1* [vision], *BBS9* [vision], *VTN* [blood coagulation and wound healing], *AKAP1* [blood coagulation], *PDE9A* [blood coagulation], *TTN* [blood coagulation], *KLK8* [response to wounding], and *PINK1* [toxic substance and hypoxia]), development (*ERBB2* [heart and peripheral nervous system], *RCAN1* [central nervous system], *AHI1* [central nervous system], *CRISPLD2* [face and lung], *KLK8* [skin], and *FRAS1* [skin]), metabolism (*NUP210*, *LDHD, ACAD9*, *TNXB, SARM1, OSBPL3* and *GBP6*), aging (*DDC8*), and immunity (*NUP210, ERBB2, HPR, HP, CTBP1, PSG3, CHMP4C, VTN, SARM1,* and *GBP6*).

In summary, we reported that Denisovan introgressions introduced 121 missense alleles into PNG and Australians, and further suggest that the 121 missense DIAs are located in evolutionarily important genes.

## Discussion

In this study, we proposed a two-phase method in identifying Denisovan introgression in PNG and Australians. We then evaluated the performance of this method by simulations. This two-phase method could effectively detect Denisovan introgressions (detection power = 79.74%), while maintaining a low false positive rate (2.44%). We applied this two-phase method to twenty PNG and Australian genomes, and identified ∼ 1.34 Gb of Denisovan introgressive haplotypes. We further found that Denisovan introgressions in 78 DIRs were under positive selection, and 121 DIAs are missense alleles. In addition, genes located in the 78 positively selected DIRs and harboring the 121 missense DIAs are of evolutionary importance. Taken together, the above findings suggest that Denisovan introgressions are evolutionarily important, and may play an important role in the evolution and local adaptation of PNG and Australians.

### Control for False Positives

In this study, we proposed a new method to identify Denisovan introgressions in non-African genomes. In this method, we carefully controlled for false positives by implementing a three-step filtering process (Ding et al. 2014b) after the identification process.

In the first (i.e., identification) phase of the two-phase method in identifying Denisovan introgressions, we identified candidates of Denisovan introgressive haplotypes based on their high divergence with Africans. However, some non-African haplotypes that are not of Denisovan origin may also have high divergence with Africans, such as haplotypes from incomplete lineage sorting (ancient population structure), background selection, and GC-biased gene conversion.

To reduce false positives, we implemented a filtering process after the identification phase. The three criteria used in the filtering process are from an earlier study (Ding et al. 2014b), and are summarized from several studies (Mendez et al. 2012a, 2012b, Ding et al. 2014a). The three criteria could effectively distinguish true Denisovan introgressions from false positives. In specific, the first criterion (close phylogenetic relationship with Denisovan) can filter out false positives from background selection. The second criterion (divergence time with Denisovan should postdate Denisovan – African divergence time) can filter out false positives from incomplete lineage sorting, background selection, and biased gene conversion. To minimize the impact of local mutation rate variation on time estimations (such as accelerated mutation rate caused by GC-biased gene conversion, Galtier and Duret 2007), we used local mutation rates calibrated from human – chimpanzee pairwise alignment in time estimations. The third criterion (rejection of alternative model by genetic length) can filter out false positives from incomplete lineage sorting. In summary, the filtering phase ensures that most false positives picked up in the identification phase were excluded from the final result. This is consistent with the simulation results (false positive rate 2.44% for PNG, and 6.44% for Eurasians, see next section). This performance, based on our simulations, is comparable to, if not better than, previously proposed approaches.

In this study, we also proposed a method to identify DIAs. The aforementioned three-step filtering process was also used in this method to filter out false positives. Specifically, for a given candidate DIA, it will be considered as a DIA only if *all* haplotypes carrying the allele satisfied the aforementioned three criteria (i.e., of Denisovan origin). False positives are not likely to pass this filter, since haplotypes carrying the false positives are not all of Denisovan origin. This stringent filtering process ensures that false positives are minimized in the final result.

In summary, we used a previously proposed three-step approach to control for false positives in this study. Since false positives are well controlled, we are confident that the conclusions of this study are not likely be influenced by false positives.

### Comparison with Other Methods

Before this study, two methods were proposed to identify Neanderthal introgressions in Eurasian genomes (Plagnol and Wall 2006, Sankararaman et al. 2014, Vernot and Akey 2014). We compared our two-phase method in identifying Denisovan introgressions in non-Africans proposed in this paper with the two other methods.

The method proposed by Vernot and Akey (2014) aims to detect the pattern of high LD between variants absent from Africans (also see Plagnol and Wall 2006, Wall et al. 2009). Despite both our method and the method of Vernot and Akey (2014) are based on the high divergence between introgressive haplotypes and Africans, we do not investigate LD of neighboring E-alleles (i.e., variants absent from Africans). Instead, we use high E-allele rate as criterion to identify introgressive haplotypes in the first phase of our method. Since the method proposed here is not based on LD, the identification of an introgressive haplotype in a given non-African genome is independent from the allele patterns in other non-African genomes.

The method proposed by Sankararaman et al. (2014) focus on alleles that are derived, not observed in Africans and consistent with Neanderthal, which is an effective way to reduce false positives. In the first phase of our method, however, we do not require E-alleles to be derived or consistent with Denisovan. Instead, we implemented a separate filtering process (i.e., the second phase) to reduce false positives. While both methods performed well in false positive control (details see below), the method proposed here may have the potential to detect introgressive haplotypes from unknown archaic hominins in future studies, since the first phase of this method is independent from the genomes of known archaic hominins.

In addition to above comparisons, we also compared the performance of our method with the two other methods. Since the evaluation of performance of our method in Results is based on the demography of PNG (detection power = 79.74%, false positive rate = 2.44%), we re-evaluated the performance of our method using the Eurasian demography here.

Demographic parameters of Eurasians and archaic hominin gene flow rate we used here are same as Vernot and Akey (2014). There are only a few small differences between our simulation and the simulation in Vernot and Akey (2014). Length of simulated Eurasian sequences is 1 Mb in our simulation, and is 50 Kb in Vernot and Akey (2014). Furthermore, our definition of archaic introgressions on the simulated sequences is: (1) coalesced first with archaic hominin (based on trees generated by msHOT); (2) diverged with archaic hominin < 550 KYA (Prüfer et al. 2014); and (3) length of segments are consistent with an introgression time after 550 KYA. In Vernot and Akey (2014), archaic introgressions were identified in trees (generated by ms) in which archaic sequence joins the tree before the simulated join time (< 700 or 400 KYA).

For simulated Eurasian genomes, the detection power of our method reached maximum of 85.31% at λ=10, with false positive rate of 6.44% (Table 1, Supplemental Table 2). The detection power and false positive rate reported in Vernot and Akey (2014) are ∼ 30% and 20%, respectively. The detection power and false positive rate reported in Sankararaman et al. (2014) are 38.4% to 55.2% and < 10%, respectively. It could therefore be suggested that the performance of our method is comparable to, if not better than, the two other methods. However, this comparison is preliminary, since simulation parameters and definition of true archaic introgressions on simulated sequences in the three studies are not exactly identical.

### Prospects

In this paper, we proposed a two-phase method in identifying Denisovan introgressions in non-Africans. This method may be applicable to other populations and other types of archaic introgressions. In combination with the method in identifying archaic introgressive alleles (i.e., DIA in this study) proposed in this study, it would be interesting to explore the functional and evolutionary importance of Neanderthal introgressions in Eurasians. Further, our two-phase method may also be applicable in identifying introgressions from unknown archaic hominins, since the first phase of our method is independent from Neanderthal and Denisovan genomes. Without the existence of known archaic hominin genome, however, extra caution should be excised to distinguish true archaic introgressions (from unknown archaic hominins) from false positives.

## Material and Methods

### The Two-phase Method in Identifying Denisovan Introgressions

In this paper, we proposed a two-phase method to identify Denisovan introgressions in non-Africans. In the first (i.e., identification) phase, we identified candidate Denisovan introgressions based on their high E-allele rate. In the second (i.e., filtering) phase, we employed a three-step filtering process to reduce false positives. Here we describe the details of this two-phase method.

In the first phase, we define *E-allele* as follows: for two alleles on a bi-allelic SNP, if an allele is absent from all African samples (507 Africans in the 1000 Genomes Project Phase 3 [1000 Genomes Project Consortium 2012], and 13 Africans from Prüfer et al. 2014, for details see below), but could be observed in PNG or Australian genome, this allele is defined as an E-allele. E-allele rate of a given segment is calculated as the number of E-alleles on the segment divided by the number of polymorphic sites in the genomic region of the segment. Simulations suggest that introgressive haplotypes have higher E-allele rate than the rest of the non-African genomes (Supplemental Text). We thus developed a hidden Markov model (HMM)-based approach with two hidden states (i.e., lower and higher E-allele rate) to partition each PNG and Australian genome into segments with different E-allele rates. Segments with *lower* E-allele rate were labeled as the *first* state, and segments with *higher* E-allele rate were labeled as the *second* state. To reduce noises, segments with length < 10 Kb were discarded. Segments with higher E-allele rates (i.e., labeled as the second state) are more likely of Denisovan origin, and will be subjected to filtering by the second phase. Below we describe the HMM-based approach.

For a non-African chromosome, we assumed it contains *M* alleles on *M* SNPs. *F*_*m*_ is the E-allele status indicator for the *m*^th^ allele. *F*_*m*_=1 indicates that this allele is an E-allele, while *F*_*m*_=0 indicates that the allele is not an E-allele. We therefore transformed *M* alleles into a string of 0 and 1s. The goal of the HMM-based approach is to partition the string into *N* segments and label each segment with a hidden state. We solved the partitioning and labeling problem with a classic Viterbi algorithm. The steps of the HMM-based approach are as follows: (1) set a penalty for state transitions (λ) and an initial E-allele rate for each hidden state; (2) use the Viterbi algorithm to partition the non-African chromosome and label each segment with a hidden state; (3) re-estimate the E-allele rate for both hidden states; (4) repeat steps 2 and 3 until convergence. We set the initial E-allele rate of the first state as the observed genome-wide E-allele rate of the non-African genomes, and the initial E-allele rate of the second state as the observed genome-wide E-allele rate of the Denisovan genome. In our search for Denisovan introgressions in PNG and Australians, the initial E-allele rates for the first and the second states were set as 0.0058 and 0.0276, respectively.

Above we described the details of the first phase of the two-phase method. Here we describe the details of the second (i.e., filtering) phase. We intended to reduce false positive rate by implementing this filtering process. Segments identified in the first phase that have higher E-allele rate (i.e., labeled as the second state) are considered as candidates of introgressive haplotypes, and are subjected to this filtering. Candidates will be considered as true Denisovan introgressive haplotypes if they satisfy all three following criteria: (1) phylogenetically closest to Denisovan, among Denisovan, Neanderthal, African, and chimpanzee sequences; (2) diverged with Denisovan < 550 KYA, and the divergence time with Denisovan is less than the divergence time with Neanderthal; and (3) genetic length > 0.01089 cM.

For the first criterion, the phylogenetic trees are reconstructed using the parsimony method implemented in PHYLIP (Felsenstein 1989). For the second criterion, we estimated the divergence time between the candidates and Denisovan, Neanderthal and African (a San individual, HGDP01029) using a maximum likelihood-based method (Mendez et al. 2012a). The mutation rates used in the time estimations are calibrated using the human (GRCh37) – chimpanzee (PanTro3) pairwise alignment, assuming a human – chimpanzee divergence time of 6,500 KYA. For the third criterion, the purpose is to reject the incomplete lineage sorting model. Based on the incomplete lineage sorting model, the candidates existed in the human gene pool before the Denisovan – African population divergence (i.e., 550 KYA). Probability of persistence of a haplotype over *T*_*p*_ years could be expressed as *p=*e^-(*θ*×*T*_*p*_)/(100×*T*_*g*_)^, where *T*_*g*_ is time per generation (20 years), and *θ* is the genetic length of the haplotype. When *p ≤* 0.05, *T*_*p*_ = 550,000 (based on the incomplete lineage sorting model), and *T*_*g*_=20, the *θ* could be computed as ≥ 0.01089 cM. Thus, the third criterion requires the candidates to be longer than 0.01089 cM to reject the incomplete lineage sorting model. Genetic map used in this study is from Hinch et al. (2011).

### Evaluation of Performance of the Two-phase Method by Simulations

In Results, we evaluated the detection power and false positive rate of our two-phase method by simulation, based on the demographic parameters of PNG. In Discussion, in order to compare with two other published methods, we evaluated the detection power and false positive rate of our method using the demographic parameters of Eurasians. Here we describe the details of the simulations.

Using msHOT (Hellenthal and Stephens 2007), we simulated non-African genomes with archaic introgressions. Demographic parameters for Africans, Eurasians, and archaic hominins are from Vernot and Akey (2014), Schaffner et al. (2005), Tennessen et al. (2012), and Gravel et al. (2011). Demographic parameters for PNG are from Prüfer et al. (2014). In specific, the demographic parameters we used are as follows. Generation time was set to 20 years, and mutation rate was set to 2×10^-8^ per base per generation (Fu et al. 2014). Population size (*N*_*e*_) of archaic hominin after their divergence with Africans is 1,500. Before 148 KYA, *N*_*e*_ of Africans was 7,300, and after 148 KYA, *N*_*e*_ of Africans was 14,474. There was exponential growth in Africans from 5,115 years ago, and present *N*_*e*_ of Africans is 424,000. Ancestors of non-Africans diverged with Africans at 60 KYA. Divergence time between Europeans and East Asians was randomly set between 36 KYA and 50 KYA. Before the European – East Asian divergence, *N*_*e*_ of Eurasians was set to the same as *N*_*e*_ of Europeans. Before 23 KYA, total *N*_*e*_ of East Asians and Europeans was randomly set between 4,000 and 18,353, with ratio of *N*_*e*_ of Europeans to *N*_*e*_ of East Asians randomly set between 1 and 2.5. We require *N*_*e*_ of Europeans less than 9,475 and *N*_*e*_ of East Asians less than 8,879 before 23 KYA. There was exponential growth in Europeans and East Asians from 23 KYA to 5,115 years ago, with *N*_*e*_ of Europeans and East Asians increased to 9,475 and 8,879, respectively. Since 5,115 years ago, there was rapid exponential growth of *N*_*e*_ in Europeans and East Asians, and the present *N*_*e*_ of Europeans and East Asians are 512,000 and 1,370,990, respectively. For Papuans, they diverged with Eurasians at 36 KYA, and we assume that their demographic history is same as East Asians before 5,115 years ago. Since 5,115 years ago, there was exponential growth in Papuans, and the present *N*_*e*_ of Papuans is 424,000 (close to Africans, Prüfer et al. 2014). Before the Eurasian divergence, migration rate between Africans and Eurasians was 1.498975×10^-4^, i.e., 1.498975×10^-4^ of Africans were new migrants from Eurasians each generation. After the Eurasian divergence, the African – European, African – East Asian, and European – East Asian migration rates were 2.498291×10^-5^, 7.794668×10^-6^, and 3.107874×10^-5^, respectively. Time of archaic hominin introgression into the ancestors of Eurasians was randomly set between 55 KYA and the Eurasian divergence time, with migration rate 0.0015. This introgression continued for 500 years. Time of archaic hominin introgression into East Asians was 500 years after the Eurasian divergence, with migration rate randomly set between 0.00015 and 0.0005. This introgression continued for 500 years. Time of archaic introgression into Papuans was 500 years after the Papuan – East Asian divergence, with migration rate of 0.0015.

In our simulation, sequence length of simulated non-African genome was set to 1 Mb. Parameters for recombination hotspot and recombination rate were from Hellenthal et al. (2008). Background recombination rate was 2.325×10^-9^ between two bases per generation. Number of hotspots for each sequence was randomly drawn from a Poisson distribution with mean value 25 (i.e., one hotspot per 40 Kb on average). Width of each hotspot was randomly set from 1 to 2 Kb. For intensity of each hotspot (*i*), we set log_10_(*i*) as random value between 1 and 2.5.

We performed simulations using the aforementioned parameters. In each simulation, we simulated 1,040 Africans, one archaic hominin, one East Asian, and one PNG (or European, for the simulation using Eurasian demography). On the simulated non-African sequences, we define simulated archaic introgressions as follows, as per the definition of true archaic introgressions: (1) they coalesced first with archaic hominin from trees generated by msHOT in simulation; (2) they diverged with archaic hominin later than 550 KYA, based on divergence times provided by msHOT; and (3) length of the segments is consistent with an introgression time after 550 KYA (i.e., longer than 0.01089 cM, see above). We repeated the simulation for 1,000 times using the PNG demography and the Eurasian demography, respectively.

We then employed our two-phase method to the simulated non-African genomes, with different λ values. The initial E-allele rates for the first and second states are the averaged E-allele rates for the simulated non-African (i.e., PNG or Eurasian) and the simulated archaic hominin sequences in the 1,000 simulations, respectively. For the simulated PNG genomes, initial E-allele rates for the first and the second states were 0.0058 and 0.0654, respectively. For the simulated Eurasian genomes, initial E-allele rates for the first and second states were 0.0044 and 0.0547, respectively. Detection power is defined as the proportion of the simulated archaic introgressions that were detected by the two-phase method, and false positive rate is defined as the proportion of the segments identified by the two-phase method that are not simulated archaic introgressions. We computed detection power and false positive rate for each integer λ value from 1 to 25 for the simulated PNG dataset, and from 1 to 15 for the simulated Eurasian dataset.

### Data Sources

After we demonstrated that our two-phase method could effectively detect Denisovan introgressions in PNG by simulation and obtained an optimized λ value, we applied this method to the sixteen PNG genomes and the four Australian genomes in the Simons Genome Diversity Project dataset.

When identifying E-alleles, 507 African genomes in the five African populations in the 1000 Genomes Project Phase 3 dataset (Esan in Nigeria, Gambian in Western Divisions in the Gambia, Mende in Sierra Leone, Luhya in Webuya, Kenya, and Yoruba in Ibadan, Nigeria, 1000 Genomes Project Consortium 2012) and 13 high-coverage African genomes from Prüfer et al. (2014, 5 Yoruba, 2 Mbuti, 2 Dinka, 2 San, 2 Mandenka) were used as Africans. In addition, we obtained high-coverage Neanderthal genome from Prüfer et al. (2014), and high-coverage Denisovan genome from Meyer et al. (2012). We phased the 13 high-coverage African genomes, the Neanderthal genome, and the Denisovan genome using the SHAPEIT software (Delaneau et al. 2013). In all of our analyses, we focused on bi-allelic SNPs, and discarded tri-allelic SNPs and insertions/deletions.

### Identifying DIAs

To explore the evolutionary importance of the identified Denisovan introgressions, we identified alleles that were introduced by Denisovan introgressions, hereby denoted as *Denisovan introgressive alleles (DIAs)*. To identify DIAs, alleles that absent in Africans but carried by all Denisovan introgressive haplotypes were considered as candidate DIAs. We then applied a stringent filter identical to the three-step approach used in the second phase of the two-phase method to control for false positives. For a given candidate DIA, it will be considered as a DIA if *all* haplotypes carrying the allele are of Denisovan origin, i.e., satisfy the following three criteria: (1) phylogenetically closest to Denisovan, among PNG and Australian haplotypes, Neanderthal, Denisovan, chimpanzee, and African (HGDP01029, a San individual); (2) diverged with Denisovan no earlier than 550 KYA; and (3) genetic length < 0.01089 cM. For the first criterion, the phylogenies are reconstructed using the neighbor-joining method implemented in PHYLIP and MEGA (Tamura et al. 2013). For the second criterion, the divergence times are estimated using the method in endez et al. (2012a). For the third criterion, the purpose is to reject the incomplete lineage sorting model.

### Scan for Positive Selection, Annotation of DIAs and Genes

We computed the standardized integrated haplotype score (iHS) for SNPs with minor allele frequency more than 5% in the twenty PNG and Australian genomes using the Whole-Genome Homozygosity Analysis and Mapping Machina (Voight et al. 2006). The genetic distance is from Hinch et al. (2011). For each DIR, the Denisovan introgression in the DIR will be considered as under positive selection if more than one (not including one) of the SNPs in strong LD (*r*^*2*^≥0.80) with the tag DIA of the DIR has standardized |iHS|≥2.00, since extreme iHS values (i.e., |iHS|≥2.00) caused by positive selection are usually in strong LD (Voight et al. 2006). We further applied the EHH test (Sabeti et al. 2002) and drew haplotype bifurcation graph (Gautier and Vitalis, 2012) for the SNP with highest |iHS| to validate that the Denisovan introgression is the target of positive selection.

We obtained annotation for the 38,877 DIAs from the Variant Effect Predictor of the Ensembl Genome Browser (Flicek et al. 2014).

## Data Access

Denisovan introgressive haplotypes in PNG and Australians identified in this study are included as Supplemental File 1. DIAs identified in this study are included as Supplemental File 2.

## Acknowledgements

This research was supported by the National Science Foundation of China (31271338 and 31330038) and the Chinese National Basic Research Program (2012CB944600).

